# Removal of disulfide from acid stress chaperone HdeA does not wholly eliminate structure or function at low pH

**DOI:** 10.1101/2021.01.17.427034

**Authors:** M. Imex Aguirre-Cardenas, Dane H. Geddes-Buehre, Karin A. Crowhurst

## Abstract

HdeA is an acid-stress chaperone that operates in the periplasm of various strains of pathogenic gram-negative bacteria. Its primary function is to prevent irreversible aggregation of other periplasmic proteins when the bacteria enter the acidic environment of the stomach after contaminated food is ingested; its role is therefore to help the bacteria survive long enough to enter and infect the intestines. The mechanism of operation of HdeA is unusual in that this helical homodimer is inactive when folded at neutral pH but becomes activated at low pH after the dimer dissociates and becomes partially unfolded. Studies with chemical reducing agents have previously suggested that the intramolecular disulfide bond of HdeA aids in the maintenance of residual structure at low pH; it is believed that this residual structure is important for clustering exposed hydrophobic residues together for the purpose of binding unfolded client proteins. In order to explore its role in HdeA structure and chaperone function we performed a conservative cysteine to serine mutation of the disulfide. We found that, although residual structure is greatly diminished at pH 2 without the disulfide, it is not completely lost; conversely, the mutant is almost completely random coil at pH 6. Aggregation assays showed that mutated HdeA, although less successful as a chaperone than wild type, still maintains a surprising level of function. These studies highlight that we still have much to learn about the factors that stabilize residual structure at low pH and the role of disulfide bonds.

## Introduction

HdeA is an ATP-independent chaperone protein [2] found in the periplasm of several pathogenic bacteria including *Shigella flexneri, Escherichia coli* and *Brucella abortus* [3-5]. It, along with sister protein HdeB, helps to prevent the irreversible aggregation of other periplasmic proteins when the organism encounters a low pH environment (found in the stomach). In this way, HdeA aids in the survival of these bacteria so that they can enter the intestines of the host and cause dysentery, a disease which affects at least 120 million people each year [6, 7]. Folded HdeA is a helical homodimer (**Figure 1**). One of its particularly unusual characteristics is that its folded state (at near-neutral pH) is its inactive state; once the bacteria enter a low pH environment (below approximately pH 3) HdeA unfolds and assumes its active role as a chaperone [3, 8].

**Figure 1.**
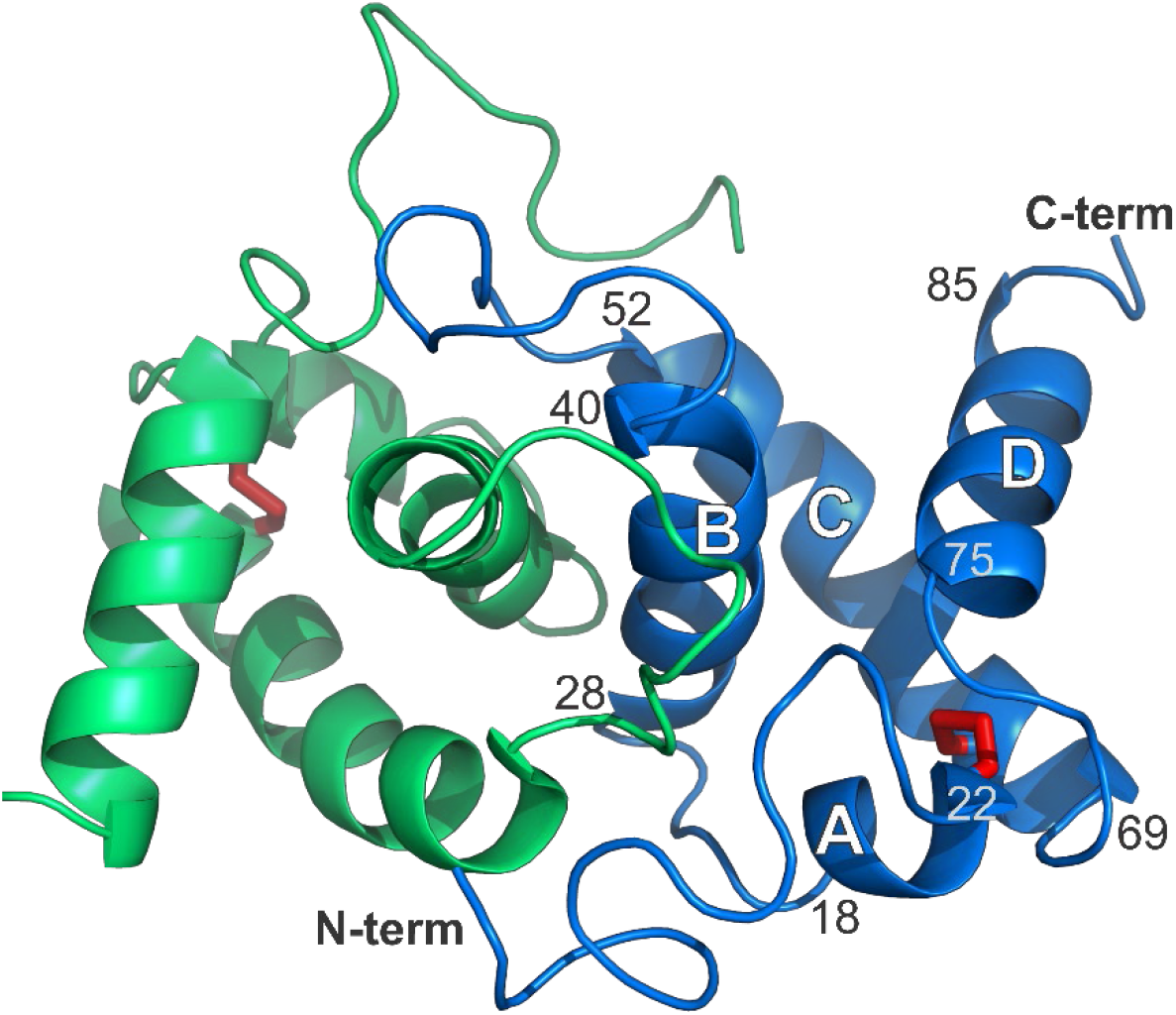
Labeled structure of the folded HdeA homodimer (PDB ID 5WYO) [2]. The blue monomer shows the locations of helices A – D (including residue number ranges) and the N-and C-termini. The disulfide bond between residues 18 and 66 is shown in red.

HdeA contains one intramolecular disulfide bond between residues 18 and 66. It has been conjectured that it may help to bring two hydrophobic sites into close contact at low pH, thereby facilitating chaperone activity by enabling the formation of an extended client binding site [9]. Although its importance is mentioned in several publications on HdeA [8, 10-12], relatively little investigation has been done on the C18-C66 disulfide bond. Hong et al. [8] originally reported that the chaperone activity of HdeA was unaffected by a reduction of the disulfide bond at low pH, but because they utilized dithiothreitol (DTT), which is functional only in the pH range 6.5 – 9.0, these observations were not valid. Tapley et al. [9] apparently demonstrated via light-scattering studies at 320 nm that, at low pH, reduced HdeA cannot act as a chaperone, although there are questions about those studies, given that DTT was once again used as the reducing agent. Finally, Zhai et al. [13] used TCEP (tris(2-carboxyethyl)phosphine), which is active over a wide pH range (1.5 – 8.5), to evaluate the chaperone activity of HdeA with a reduced disulfide at low pH, using SDS-PAGE to detect client protein aggregates. The authors suggest that the chaperone activity of reduced HdeA is heavily compromised, although their gel seems to show almost equal quantities of soluble and precipitated client protein in the presence of TCEP-treated HdeA [13]. In addition, it is unclear whether HdeA remains reduced during the assay, since the authors describe a ten-fold dilution of the TCEP at one stage.

Considering that researchers assert the presence of the disulfide is crucial for HdeA chaperone activity but have not demonstrated this definitively, we decided to prepare a double mutant of HdeA in which the cysteines were conservatively mutated to serines. These mutations eliminate the possibility of disulfide bond formation without eliminating side chain hydrogen bond capabilities and allow for studies in the absence of chemical reductants. Our results show that the loss of the disulfide bond eliminates the ability of HdeA-C18S-C66S (also called C18S-C66S) to fold at neutral pH. The mutant has near-random coil structure at pH 6.0, but surprisingly *gains* some secondary structure content at low pH, although it is still lower than that of wild type HdeA at pH 2.0. Additionally, a comparison of chaperone activity of C18S-C66S versus TCEP-reduced wild type on client protein malate dehydrogenase (MDH) indicates that both have notable (although not complete) success in keeping the client protein soluble.

## Materials and Methods

Isotopes were obtained from Cambridge Isotope Laboratories, and chromatography columns were from GE Lifesciences.

### Preparation of HdeA

Site-directed mutagenesis was executed using the QuikChange Lightning kit from Agilent. HdeA-C18S-C66S was expressed and purified as outlined previously for wild type HdeA [11, 14, 15], with the exception that a Superdex 75 HiLoad 26/600 column was required rather than the usual HR 10/30 column. All samples were uniformly ^13^C/^15^N labeled.

### NMR experiments

C18S-C66S samples were prepared for NMR through dialysis into 50 mM citrate buffer at the desired pH and had final concentrations in the range of 0.1 – 1.0 mM. Since no chemical shift changes were observed as a function of sample concentration, we were not concerned about the impact of varied protein concentration on the results presented. Reduced samples of wild type HdeA contained 5 mM tris(2-carboxyethyl)phosphine (TCEP); in order to obtain spectra at pH 6.0 the protein sample was first reduced in the unfolded state at low pH and then dialyzed to 6.0 in the presence of TCEP. NMR data were obtained at 25 °C on an Agilent DD2 600 MHz spectrometer equipped with a triple resonance probe. All raw data were processed using NMRPipe/NMRDraw [16, 17] and the resulting spectra were viewed and analyzed using NMRViewJ [18, 19].

#### Chemical shift assignment

Because C18S-C66S is unfolded at pH 6.0 and 2.0, backbone chemical shift assignments at both pHs required HNCaCb, CbCa(CO)NH and HNN [20] experiments; additionally, HNCO and HN(Ca)CO experiments were required at pH 6.0. The assignment data have been deposited at the BioMagResBank (BMRB), acquisition number 50437. Wild type HdeA assignments at pH 2.0 (BMRB acquisition number 50421, [1]) and pH 6.0 (BMRB acquisition number 19165, [14]) were reported previously, and assignments of spectra from wild type HdeA in TCEP were made by overlaying with mutant spectra.

Backbone amide chemical shift differences (CSDs, or Δδ) were calculated using the equation:

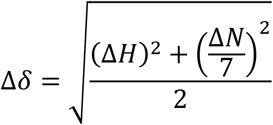

where ΔH and ΔN refer to the differences in backbone ^1^H_N_ and ^15^N chemical shifts for a given residue between different pHs or between wild type and mutant at a specific pH.

#### Secondary structure propensity

SSP analysis was performed on HdeA and HdeA-C18S-C66S using only Cα and Cβ chemical shifts, as recommended for an unfolded protein [21].

### Aggregation assays

The chaperone activity of C18S-C66S to prevent or rescue aggregation of malate dehydrogenase (MDH) was tested at pH 2.0 and 6.0, using Aggregation Buffer consisting of 20 mM citrate, 100 mM sodium chloride and 150 mM ammonium sulfate [13]. All samples contained 10 µM MDH and some contained 30 µM wild type HdeA or mutant, plus 5 mM TCEP when appropriate. At pH 2.0, the mixtures were incubated at 37 °C for one hour, then centrifuged at 14,000 xg for 10 minutes to separate the supernatant and pellet. The pellets were washed once with Aggregation Buffer and centrifuged again (to remove surface supernatant); the supernatants were partially neutralized with 0.13 volumes of 0.5 M sodium phosphate, pH 8 in advance of analysis via SDS-PAGE. At pH 6.0, MDH was pre-treated with 2 M guanidinium hydrochloride in Aggregation Buffer and incubated at 100 °C for 20 minutes to ensure MDH was denatured before mixing with HdeA (wild type or mutant), and to test the ability of the chaperone to “ rescue” aggregated MDH at pH 6.0. The mixtures were incubated at 37 °C for one hour, then centrifuged to separate the supernatant and pellet. Pellets were washed once with Aggregation Buffer and centrifuged again. All samples were run on 15% SDS-PAGE gels.

## Results and Discussion

### Chemical shift assignment

Upon overlaying the ^15^N-HSQC of C18S-C66S with wild type HdeA at pH 2.0, it was clear that the mutations significantly alter the residual low-pH structure of HdeA (**Figure S1a**). After comparing the spectra of C18S-C66S at pH 2.0 and 6.0 (**Figure S1b**) it was also clear that the loss of the disulfide bond results in unfolded protein at both pHs, but the ensembles of structures are not the same. To aid in chemical shift assignment of unfolded protein at each pH, we employed the 3D HNN experiment [20] in addition to the HNCaCb/CbCa(CO)NH and, at pH 6.0, the HN(Ca)CO/HNCO suites of experiments. In the end we achieved near-complete backbone assignment: 99% of the backbone ^1^H/^15^N atoms and 99% of the Cα/Cβ atoms at pH 2.0, as well as 95% of the backbone ^1^H/^15^N atoms, 99% of the Cα/Cβ atoms and 100% of the C(O) atoms at pH 6.0 were successfully assigned. Missing assignments at pH 2.0 were located at P12 (Cβ), G34 (Cα) and K87 (^1^H/^15^N), while at pH 6.0 missing assignments were at D2 (^1^H/^15^N), K42 (^1^H/^15^N), K44 (Cβ), D83 (^1^H/^15^N) and I85 (^1^H/^15^N). The data have been deposited at the BioMagResBank (BMRB), acquisition number 50437.

### Chemical shift differences indicate some long-range impacts of disulfide removal

The sites of the largest differences (Δδ, or CSD) in backbone amide chemical shift were evaluated. When comparing wild type HdeA to C18S-C66S at pH 2.0, it is noteworthy that the amide shifts of the first and last 15 residues in the protein are essentially indistinguishable (**Figure 2a**). As one might expect, the largest chemical shift differences can be found near (but not always at) the mutated cysteines (**Figure 2b**). In a trend that is consistent with other experiments we have done [1, 11], the largest changes are on the C-terminal side of position 18 (residues 20 – 25) and the N-terminal side of position 66 (residues 57 – 65). Within the folded structure, portions of these two regions (residues 20 – 25 and 62 – 65) are adjacent to each other; this proximity seems to be maintained in the unfolded state of the wild type, primarily due to the disulfide tether [11]. However, residues 57 – 61 extend along the entire C-terminal half of helix C, far from residues 20 – 25 and far from the site of the removed disulfide in the mutant (**Figure 2b**). When assessing secondary structure propensities (based on chemical shift assignments), the loss of the disulfide not only eliminates the helical propensity of helix C that is seen in the wild type, its secondary structure flips to weak β-sheet content (**Figure 3** and **Table S1**); this, or an allosteric effect, could explain the large chemical shift changes.

**Figure 2.**
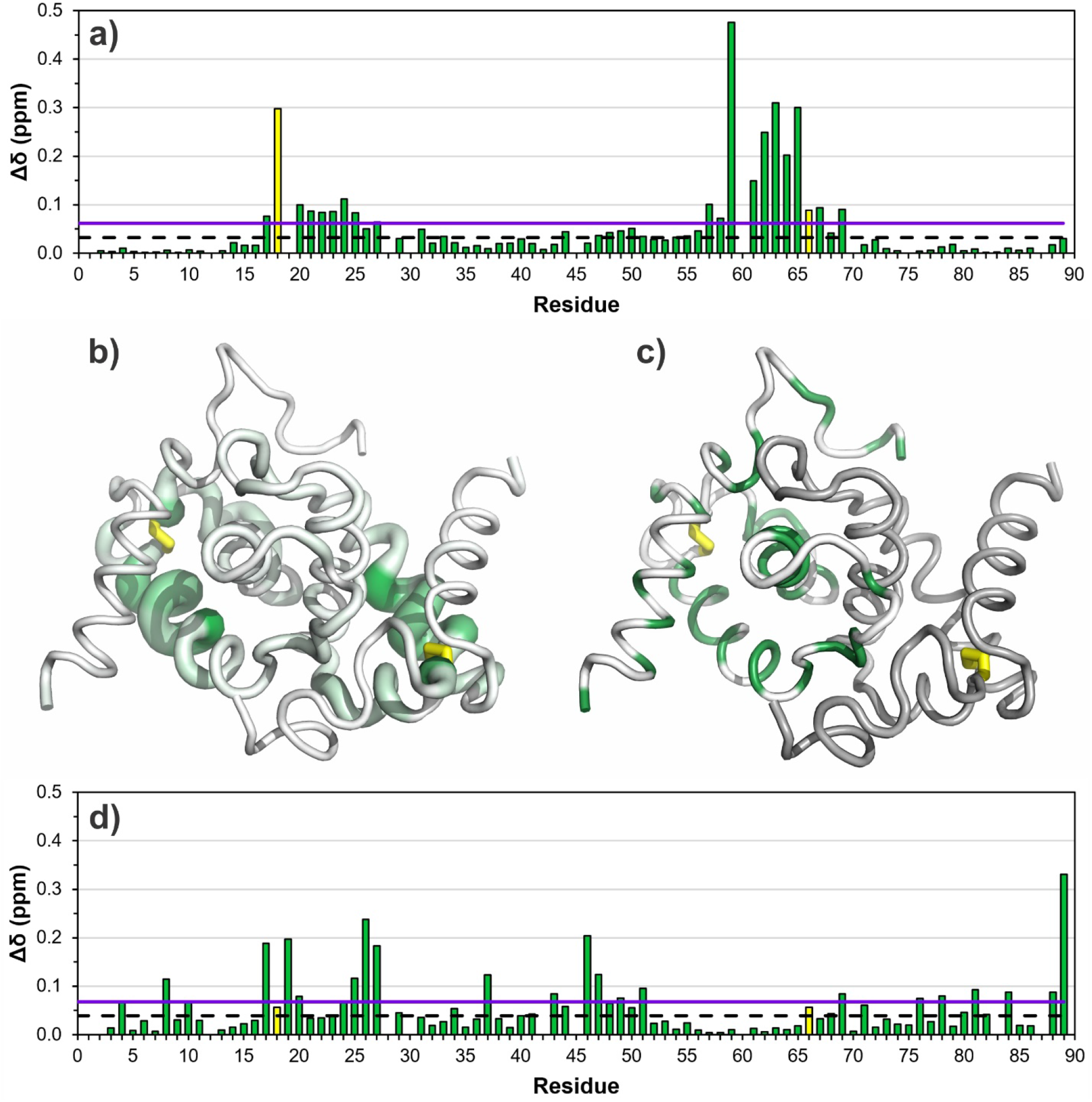
Evaluation of the amide ^1^H and ^15^N chemical shift differences (CSD, or Δδ), as a function of residue. **a)** CSD between wild type HdeA and C18S-C66S at pH 2.0. The positions of the mutation sites are colored in yellow. The black dashed horizontal line corresponds to the average Δδ (minus 10% outliers), and the purple horizontal line indicates one standard deviation above the mean. **b)** The Δδ values from **a)** are plotted on the folded structure of HdeA. Residues with higher Δδ are shown with darker green and larger cartoon radius, and the site of the wild type disulfide is colored yellow. **c)** Positions of hydrophobic groups in HdeA, colored green on one chain. **d)** CSD between C18S-C66S at pH 6.0 and 2.0.

**Figure 3.**
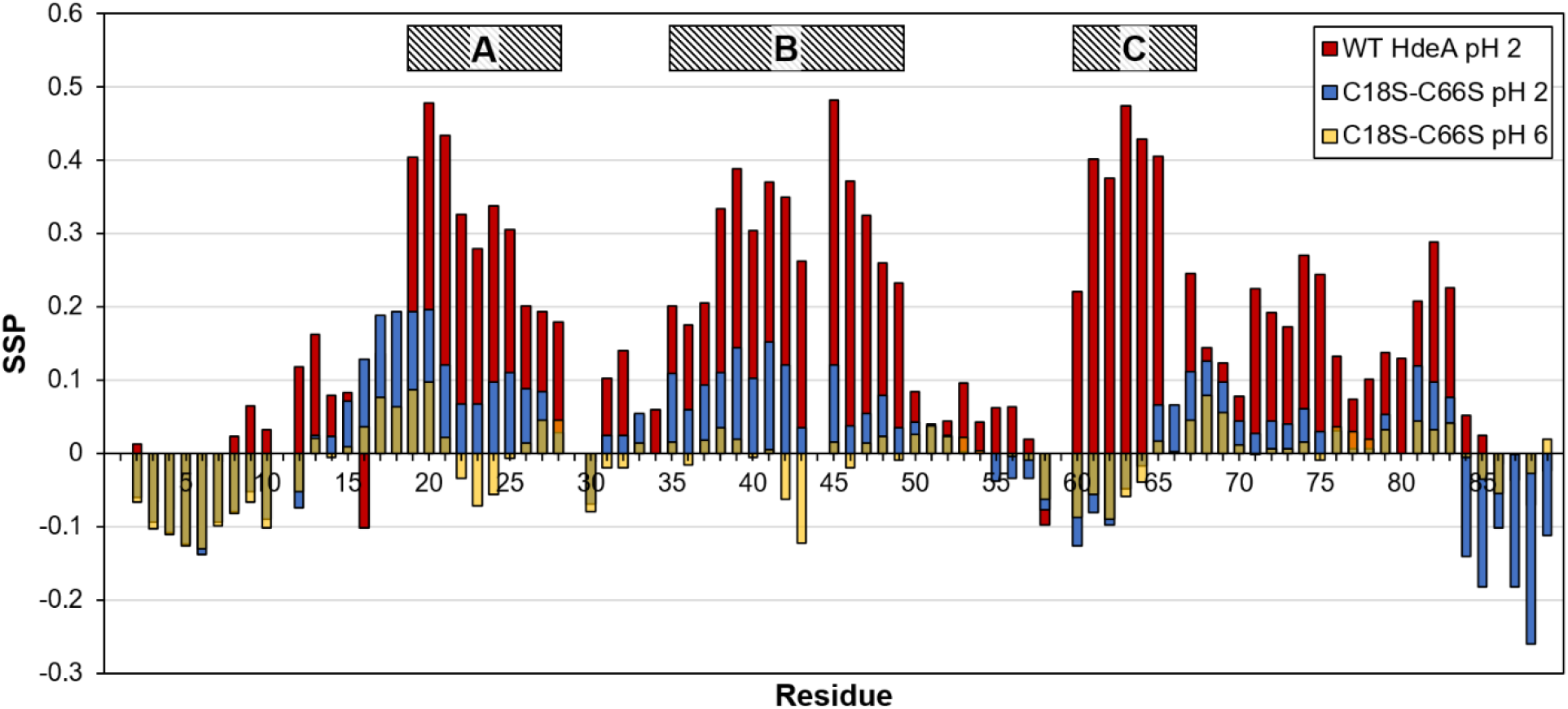
Plot of secondary structure propensities (SSP) as a function of residue number for wild type HdeA at pH 2.0 (red) and HdeA-C18S-C66S at pH 2.0 (blue) and pH 6.0 (yellow, shown as 50% transparent in order to show values from the other samples underneath). All data were calculated using only Cα and Cβ chemical shifts, as recommended for unfolded proteins. Positive SSP values indicate helical secondary structure, with maximum possible value of 1.0 (indicating 100% helical propensity) and negative SSP values indicate β-sheet secondary structure (where a value of -1 would indicate 100% β-sheet forming propensity). Approximate positions of residual helix structure in wild type HdeA at low pH are indicated at the top of the plot [1].

Like the groups described above, residues on the opposite side of each cysteine, at the N-terminal side of position 18 (including 14 – 16) and the C-terminal side of position 66 (residues 70 – 74), are also adjacent to each other in the folded protein; however, these segments of the mutant do not have significant differences in chemical shift at pH 2.0 compared to wild type. In folded wild type HdeA, this region acts like a clasp that is opened at low pH to expose hydrophobic client binding sites as part of its chaperone activation [1]. Refer to **Figure 2c** for the positions of each hydrophobic residue in HdeA. The chemical shift similarities imply structural similarities in this region between wild type and C18S-C66S at pH 2.0; these results support the notion that this region of the protein is indeed open and less structured in the wild type when the clasp is released at low pH.

Upon initial inspection, the chemical shift differences between C18S-C66S at pH 6.0 and 2.0 (**Figure 2d**), seem to have no pattern to the magnitude of Δδ value. However, upon closer inspection, it becomes clear that 14/19 residues with Δδ > 1 standard deviation above the mean are aspartate or glutamate, all of which undergo neutralization in transitioning from pH 6.0 to 2.0. The remaining residues likely have large chemical shift perturbations due to their proximity to these ionizable groups.

### Secondary structure propensity analysis shows that the mutant maintains some structure at low pH but is virtually random coil in near-neutral conditions

As mentioned briefly above, the chemical shift data were also used to calculate and compare secondary structure propensities (SSP). Using the recommendation by Marsh et al. [21], only Cα and Cβ shifts were used to calculate SSP values for the unfolded proteins. **Figure 3** shows an overlay of SSP values as a function of residue number for wild type and C18S-C66S at pH 2.0, as well as C18S-C66S at pH 6.0. See **Table S1** for the numerical values and **Figure S2** for a plot showing the change in mutant SSP (ΔSSP) between pH 2.0 and 6.0 as a function of residue number. As has been observed in previous publications [1, 12], even at pH 2.0 the “ unfolded” wild type maintains notable residual helical secondary structure, albeit in slightly shifted positions compared to the folded structure [1]. However, what is truly remarkable when evaluating the SSP values for C18S-C66S is that the mutant has more structural content at low pH compared to near-neutral conditions (**Figure 3**). At pH 6.0 C18S-C66S has extremely low SSP values, almost exclusively in the -0.1 to 0.1 range, suggesting that the protein is very close to random coil. However, at pH 2.0, the SSP values in the mutant are significantly strengthened in almost every region of the protein compared to pH 6.0; its helical secondary structure propensity even outperforms wild type at several residues on the N-terminal side of the C18S mutation. Additionally, at pH 2.0 the C18S-C66S mutant displays notable β structure propensity at the C-terminus, while wild type shows no persistent secondary structure in that region (**Figure 3** and **Table S1**). C18S-C66S also displays greater β propensity at the N-terminus at both pHs compared to unfolded wild type, suggesting that the N-and C-termini of the mutant may have an increased tendency to form semi-stable β-sheet structures with each other; this is even higher than the tendency we observed when modeling the unfolded wild type with MD simulations [11]. In trying to explain the gain of structure in the mutant at pH 2.0 compared to pH 6.0 we wondered whether there is prior experimental evidence that hydrogen bonds are generally stronger at low pHs compared to neutral pH. If this were so, we could argue that even weak hydrogen bonds between S18 and S66 (or between other residues) in the mutant could help maintain a portion of the secondary structure that is observed in the wild type at pH 2.0. Alternatively, there may be other features, in addition to the disulfide in wild type HdeA, that help the protein maintain a partially folded conformation at low pH and that are strengthened in the mutant.

Unfortunately, we have yet to find published evidence of either hypothesis; the phenomenon may be worth pursuing computationally.

### C18S-C66S has similar (but not identical) chemical shifts to wild type HdeA in TCEP

We were curious to see whether our double mutant behaved similarly to wild type HdeA in the presence of the reducing agent TCEP. When comparing the CSDs between reduced wild-type and C18S-C66S at both pH 2.0 and 6.0 (**Figures S3** and **S4**), the presence of TCEP clearly results in a similar ensemble of unfolded structures as those observed for the double mutant. The chemical shift perturbations are, overall, an order of magnitude smaller than comparisons between unfolded, oxidized wild type HdeA and C18S-C66S at pH 2.0 (compare **Figure 2a** and **S4**). Even so, the CSDs at each pH follow distinctly different trends. At pH 6.0, the CSD values between wild type in TCEP and C18S-C66S are very low, except in the region immediately surrounding the mutation sites (as one might expect). However, at pH 2.0, the average CSD is more than double what is observed at pH 6.0, and although the largest values are also near the mutation sites, there are other notably high values near the N-terminus and between residues 47 – 51 (**Figure S4a**). Neither of these regions are close to the disulfide in the folded state, but the N-terminus of one protomer is in contact with residues 47 – 51 in the other protomer of the folded dimer. In looking at the residues with the highest CSD in those two regions, five out of the six are aspartates. This suggests that TCEP may specifically interact with aspartates in solution, altering their backbone amide chemical shifts, but we cannot explain why the other aspartates (or glutamates) in the protein are not similarly affected.

### Loss of the disulfide does not wholly eliminate chaperone activity in HdeA

We were interested to evaluate the chaperone activity of C18S-C66S, and to determine whether the activity of wild type HdeA in TCEP was comparable. It has been suggested previously that the disulfide may have a crucial role in chaperone activation by clustering hydrophobic patches in HdeA and creating a larger client binding site in the partially unfolded state [9]. It was therefore expected that neither the mutant nor chemically reduced wild type HdeA would have the ability to protect a client protein from aggregation. We performed aggregation assays at pH 2.0 and 6.0, employing methods similar to those reported by other groups, using malate dehydrogenase (MDH) as a stand-in for a client protein [9, 13, 22, 23]. **Figures S5** and **S6** show the gel images and plots of quantified band density, and **Table 1** summarizes the density values. The results at pH 6.0 are as one might expect – even wild type HdeA is not expected to rescue aggregated MDH since it is folded and generally inactive as a chaperone at that pH (**Figure S5**). At pH 2.0, MDH without chaperone is found almost exclusively in the pellet and MDH in the presence of wild type HdeA is found almost exclusively in the supernatant, as expected (**Figure S6**). The results become interesting when evaluating the chaperone activity for C18S-C66S and

**Table 1.** Relative volume of the gel bands from the aggregation assays performed at pH 6.0 and 2.0.

|  | pH 6.0 | pH 2.0 |
| --- | --- | --- |
| P<sup>a</sup> – MDH only | 96.2<sup>b</sup> | 89.4 |
| S<sup>a</sup> – MDH only | 3.8 | 10.6 |
| P – WT<sup>c</sup> HdeA | 94.6 | 1.5 |
| S – WT HdeA | 5.4 | 98.5 |
| P – WT HdeA + TCEP | 91.4 | 59.8 |
| S – WT HdeA + TCEP | 8.6 | 40.2 |
| P – HdeA-C18S-C66S | 95.5 | 63.5 |
| S – HdeA-C18S-C66S | 4.5 | 36.5 |
| P – HdeA-C18S-C66S + TCEP | 95.5 | 64.9 |
| S – HdeA-C18S-C66S + TCEP | 4.5 | 35.1 |

<sup>a</sup> P and S labels represent lanes containing pellet (aggregated MDH) or supernatant (soluble MDH), respectively.
<sup>b</sup> Numbers estimated using volume integration data from Bio-Rad Image Lab software. Values correspond to % relative volume (each pellet-supernatant pair adds up to 100%).

HdeA plus TCEP; although there is a sizeable portion of insoluble client protein, at least one-third of the MDH can be found in the supernatant of each sample (**Table 1** and **Figure S6**). This implies that, while the disulfide bond is important to the success of HdeA as a chaperone, the lack of disulfide does not fully eliminate its capabilities.

Although the results show that wild type HdeA plus TCEP has similar chaperone properties as the C18S-C66S mutant, the NMR data indicate there are still differences in the conformational ensembles (**Figures S3a** and **S4a**). Given these results, we argue it is advantageous to use the mutant in any future studies, since it eliminates the variability and complication of employing a chemical additive which may also have unintended effects on other solution components, such as a client protein.

### Results provide evidence for the maintenance of some hydrophobic clustering in mutant HdeA

Considering that HdeA maintains some chaperone activity at low pH, even in the absence of the disulfide, it is worthwhile to take a deeper look at the data. We have already suggested that there may be hydrogen bonding between the serines in C18S-C66S that can help maintain some residual structure (and therefore chaperone activity) at pH 2.0, but this alone is unlikely to fully explain our assay results. Helix C in wild type HdeA contains numerous hydrophobic groups and constitutes a portion of client binding site I proposed by Yu et al. [12]. However, at pH 2.0 a large segment of this region shows significant chemical shift perturbation in C18S-C66S compared to wild type (**Figure 2a**), and most of its secondary structure propensity is lost (**Figure 3**). These data therefore suggest that there is little residual hydrophobic clustering in this region and therefore does not contribute to the protein’s chaperone activity. On the other hand, when evaluating the region corresponding to helix B and the BC loop (as seen in folded HdeA), the mutant has very similar amide chemical shifts to that of wild type at pH 2.0 (**Figure 2**), and it maintains some residual helical structure (**Figure 3**). This region is the location of the proposed client binding site II [12], and is also the same segment that is believed to become accessible (due to the opening of the clasp region) when the wild type unfolds and becomes chaperone-active [1]. In addition, the SSP data show that there is increased β structure propensity at both the N-and C-termini in C18S-C66S (**Figure 3**) compared to wild type; previous studies have provided evidence that the termini transiently form a β-sheet structure as part of HdeA’s chaperone activation [11]. Taken together, it is possible that this strengthened β-sheet formation (compared to wild type) helps C18S-C66S maintain a more compact shape, subsequently preserving the hydrophobic cluster in the client binding site near residues 28 – 39 and possibly explaining the partial chaperone activity of the mutant at pH 2.0. Future studies will investigate the importance to chaperone activity of this β-sheet formation as well as the specific location of the disulfide.

### Conclusions

Replacement of the cysteines that form the disulfide bond in HdeA with serines results in a near-random coil structure at pH 6.0, but notably higher structural content at pH 2.0. HdeA-C18S-C66S also retains greater low-pH chaperone activity than previously suggested. Both low-pH results for the mutant are unexpected, highlighting the need for more investigation of positions 18 and 66, located at a “ pinch point” of the HdeA structure [1], as well as other residues in the vicinity and at the N-and C-termini that may collectively maintain partial chaperone activity when the cysteines are removed or reduced. These results underscore the complexities of the structure-function relationship in this acid stress chaperone and may open new areas of inquiry into the role of long-range disulfide bonds in small proteins.

## Acknowledgements

We gratefully acknowledge the NIH for research support (SC3-GM116745) and the NSF for funding the purchase of our NMR spectrometer (CHE-1040134). Sincere thanks also go to Ranjith Muhandiram (University of Toronto) for assistance with the HNN assignment experiment and for helpful discussions.

