## Supplementary material for "Removal of disulfide from acid stress chaperone HdeA does not wholly eliminate structure or function at low pH"

**Table S1: Secondary structure propensity (SSP) values for wild type and mutant**

| | SSP values for: | | |
|---|---|---|---|
| Residue number | WT HdeA at pH 2.0 | HdeA-C18S-C66S at pH 2.0 | HdeA-C18S-C66S at pH 6.0 |
| 1 | | | |
| 2 | 0.013 | -0.060 | -0.066 |
| 3 | -0.066 | -0.094 | -0.103 |
| 4 | -0.077 | -0.108 | -0.110 |
| 5 | -0.051 | -0.124 | -0.126 |
| 6 | -0.018 | -0.138 | -0.130 |
| 7 | -0.012 | -0.094 | -0.099 |
| 8 | 0.023 | -0.079 | -0.082 |
| 9 | 0.065 | -0.052 | -0.067 |
| 10 | 0.032 | -0.090 | -0.101 |
| 11 | | | |
| 12 | 0.118 | -0.074 | -0.052 |
| 13 | 0.162 | 0.025 | 0.021 |
| 14 | 0.079 | 0.023 | -0.005 |
| 15 | 0.083 | 0.072 | 0.009 |
| 16 | -0.101 | 0.129 | 0.036 |
| 17 | 0.137 | 0.189 | 0.077 |
| 18 | | 0.193 | 0.064 |
| 19 | 0.404 | 0.194 | 0.087 |
| 20 | 0.478 | 0.196 | 0.098 |
| 21 | 0.434 | 0.121 | 0.022 |
| 22 | 0.326 | 0.068 | -0.034 |
| 23 | 0.280 | 0.068 | -0.072 |
| 24 | 0.338 | 0.097 | -0.056 |
| 25 | 0.306 | 0.111 | -0.007 |
| 26 | 0.202 | 0.088 | 0.014 |
| 27 | 0.194 | 0.085 | 0.046 |
| 28 | 0.179 | 0.028 | 0.046 |
| 29 | | | |
| 30 | -0.012 | -0.069 | -0.080 |
| 31 | 0.102 | 0.024 | -0.020 |
| 32 | 0.140 | 0.024 | -0.020 |
| 33 | 0.052 | 0.054 | 0.014 |
| 34 | 0.060 | | |
| 35 | 0.202 | 0.109 | 0.016 |
| 36 | 0.175 | 0.060 | -0.016 |
| 37 | 0.205 | 0.093 | 0.018 |
| 38 | 0.334 | 0.110 | 0.035 |
| 39 | 0.388 | 0.144 | 0.019 |
| 40 | 0.304 | 0.103 | -0.005 |
| 41 | 0.370 | 0.152 | 0.005 |
| 42 | 0.349 | 0.121 | -0.062 |
| 43 | 0.263 | 0.035 | -0.122 |

| | | | |
|---|---|---|---|
| 44 | | | |
| 45 | 0.482 | 0.121 | 0.015 |
| 46 | 0.372 | 0.037 | -0.019 |
| 47 | 0.325 | 0.054 | 0.014 |
| 48 | 0.260 | 0.079 | 0.023 |
| 49 | 0.232 | 0.035 | -0.009 |
| 50 | 0.085 | 0.043 | 0.026 |
| 51 | 0.036 | 0.040 | 0.037 |
| 52 | 0.044 | 0.025 | 0.023 |
| 53 | 0.096 | 0.003 | 0.022 |
| 54 | 0.043 | | 0.004 |
| 55 | 0.062 | -0.038 | 0.000 |
| 56 | 0.063 | -0.034 | -0.004 |
| 57 | 0.019 | -0.034 | -0.009 |
| 58 | -0.098 | -0.077 | -0.062 |
| 59 | | | |
| 60 | 0.221 | -0.126 | -0.087 |
| 61 | 0.402 | -0.081 | -0.056 |
| 62 | 0.375 | -0.098 | -0.090 |
| 63 | 0.474 | -0.048 | -0.058 |
| 64 | 0.429 | -0.017 | -0.039 |
| 65 | 0.405 | 0.066 | 0.017 |
| 66 | | 0.066 | 0.003 |
| 67 | 0.245 | 0.112 | 0.045 |
| 68 | 0.144 | 0.126 | 0.079 |
| 69 | 0.123 | 0.098 | 0.056 |
| 70 | 0.078 | 0.044 | 0.011 |
| 71 | 0.225 | 0.027 | -0.001 |
| 72 | 0.192 | 0.044 | 0.007 |
| 73 | 0.173 | 0.040 | 0.007 |
| 74 | 0.270 | 0.061 | 0.015 |
| 75 | 0.244 | 0.030 | -0.009 |
| 76 | 0.132 | 0.031 | 0.036 |
| 77 | 0.074 | 0.007 | 0.030 |
| 78 | 0.101 | 0.006 | 0.019 |
| 79 | 0.138 | 0.053 | 0.032 |
| 80 | 0.130 | | |
| 81 | 0.208 | 0.120 | 0.044 |
| 82 | 0.289 | 0.098 | 0.032 |
| 83 | 0.226 | 0.077 | 0.041 |
| 84 | 0.052 | -0.140 | -0.005 |
| 85 | 0.025 | -0.182 | -0.035 |
| 86 | -0.015 | -0.102 | -0.055 |
| 87 | -0.036 | -0.182 | -0.002 |
| 88 | -0.070 | -0.260 | -0.027 |
| 89 | -0.023 | -0.112 | 0.020 |

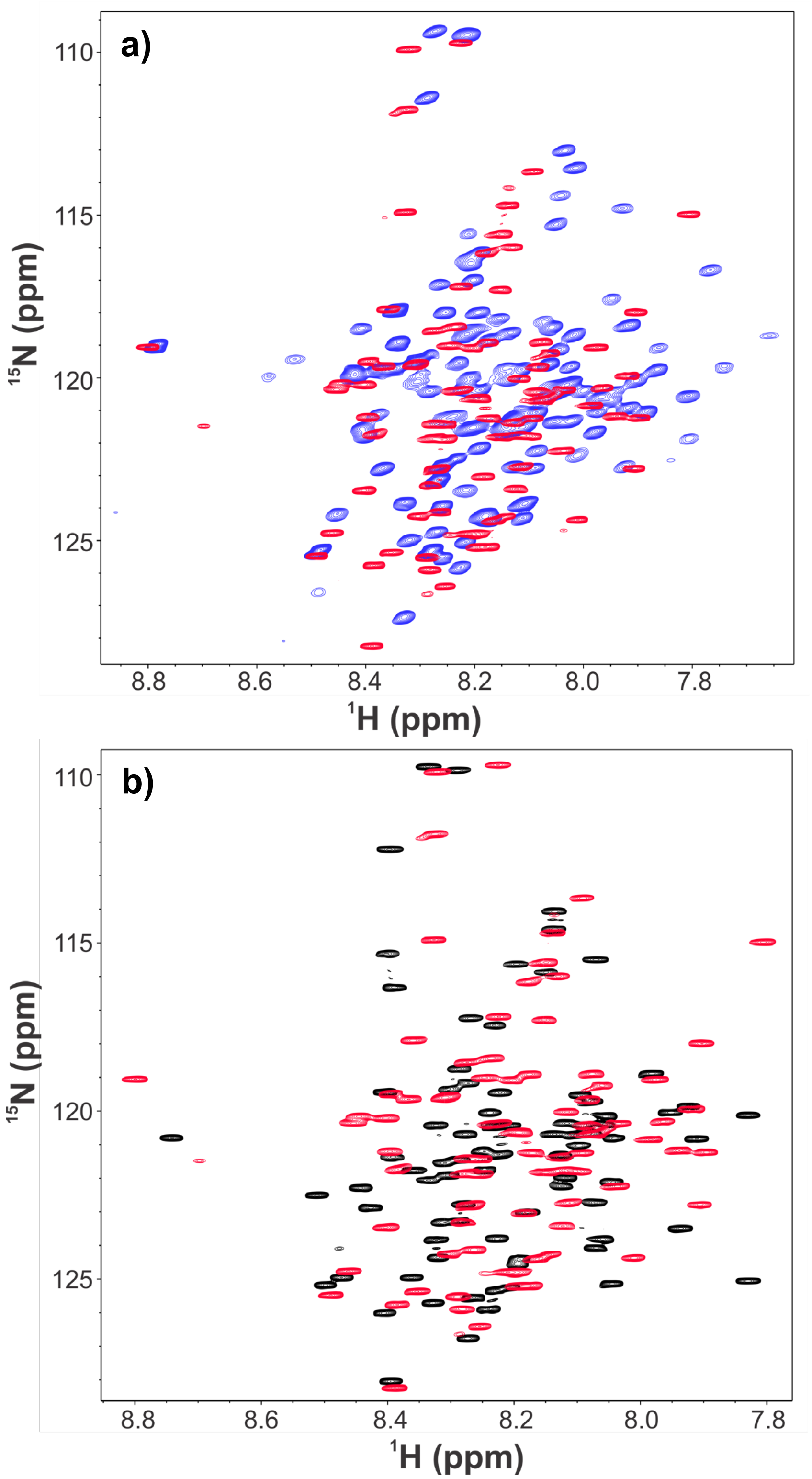


**Figure S1.** Overlays of $^{15}$N-HSQC spectra of **a)** HdeA-C18S-C66S at pH 2.0 (red) and WT HdeA at pH 2.0 (blue), **b)** HdeA-C18S-C66S at pH 6.0 (black) and pH 2.0 (red).

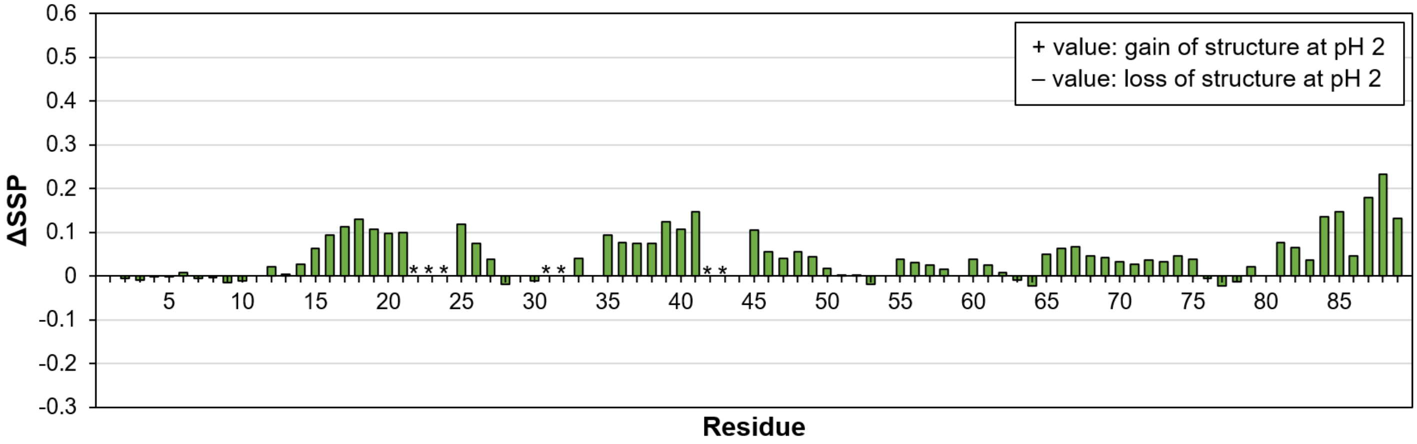


**Figure S2.** Plot of the difference in secondary structure propensity (ΔSSP) between HdeA-C18S-C66S at pH 2 versus pH 6 as a function of residue number. Any positive number (regardless of type of secondary structure) indicates a gain of structure of the mutant at pH 2 compared to pH 6, while a negative number indicates a loss of secondary structure at that position. In cases where there is a substantial change in SSP from helical to sheet or vice-versa, those positions are marked with *. For sake of consistency, the range of values on y-axis is the same as shown in Figure 3.

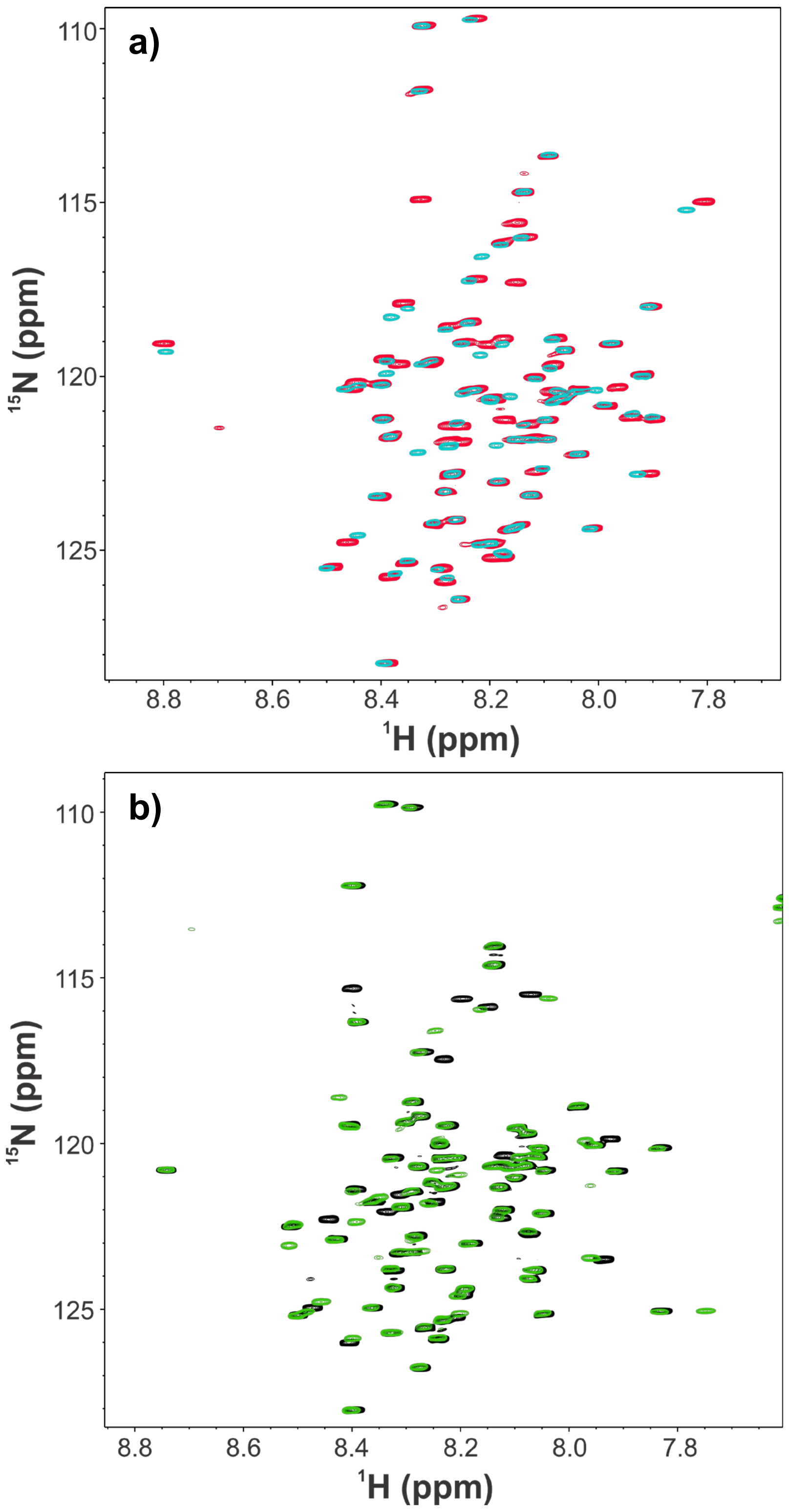


**Figure S3.** Overlays of $^{15}$N-HSQC spectra of **a)** HdeA-C18S-C66S at pH 2.0 (red) and WT HdeA treated with TCEP at pH 2.0 (cyan), **b)** HdeA-C18S-C66S at pH 6.0 (black) and WT HdeA treated with TCEP at pH 6.0 (green)

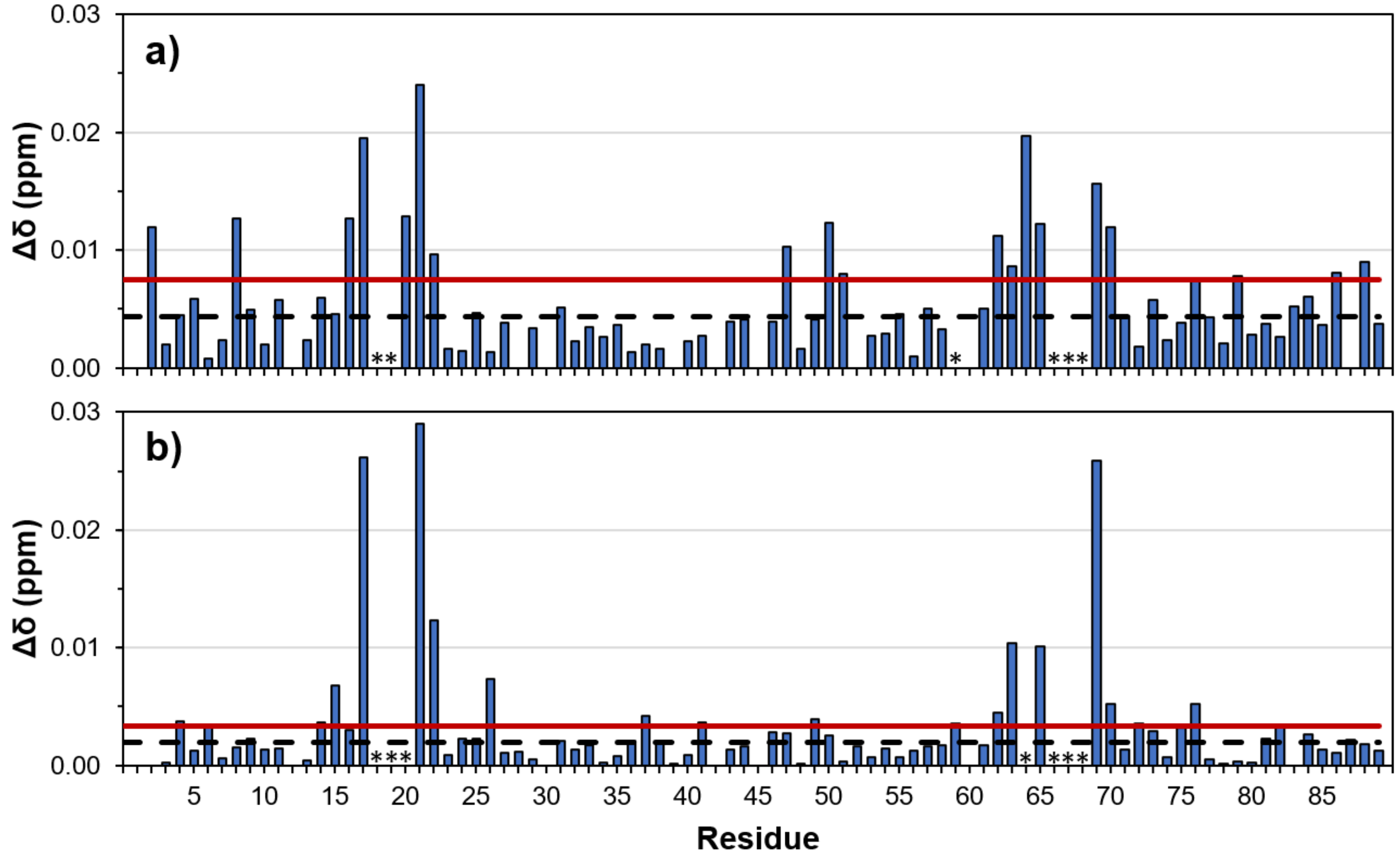


**Figure S4.** Evaluation of the amide $^{1}H$ and $^{15}N$ chemical shift differences (CSD, or Δδ), as a function of residue, between **a)** wild type HdeA in 5 mM TCEP and C18S-C66S at pH 2.0, and **b)** wild type HdeA in 5 mM TCEP and C18S-C66S at pH 6.0. The black dashed horizontal line corresponds to the average CSD (minus 10% outliers), and the red horizontal line indicates one standard deviation above the mean. Chemical shift assignments for WT in TCEP were obtained via overlay with the C18S-C66S spectrum. Residues which could not be assigned due to large chemical shift differences are indicated with a star (*).

a)

b)

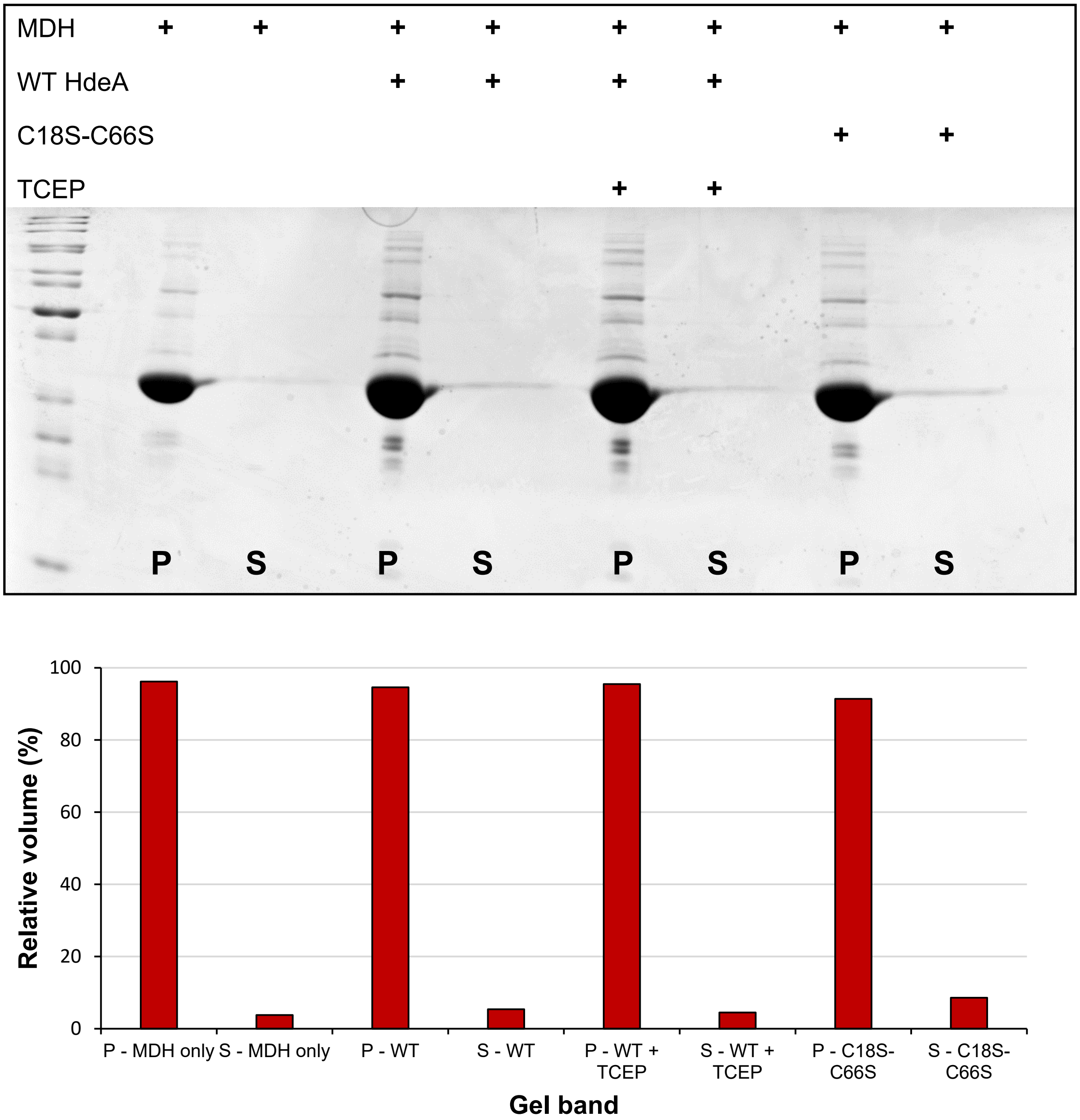


**Figure S5. a)** SDS-PAGE gel showing the HdeA chaperone activity ("rescue") assay at pH 6.0. "P" and "S" labels represent lanes containing pellet (aggregated MDH) or supernatant (soluble MDH), respectively. "+" signs in the table above the gel indicate the presence of the indicated component. Gel image has been adjusted for brightness and contrast, but the adjustment has not eliminated or obscured any bands. **b)** Plot of volume integration data from Bio-Rad Image Lab software. Values correspond to % relative volume (each pellet-supernatant pair adds up to 100%).

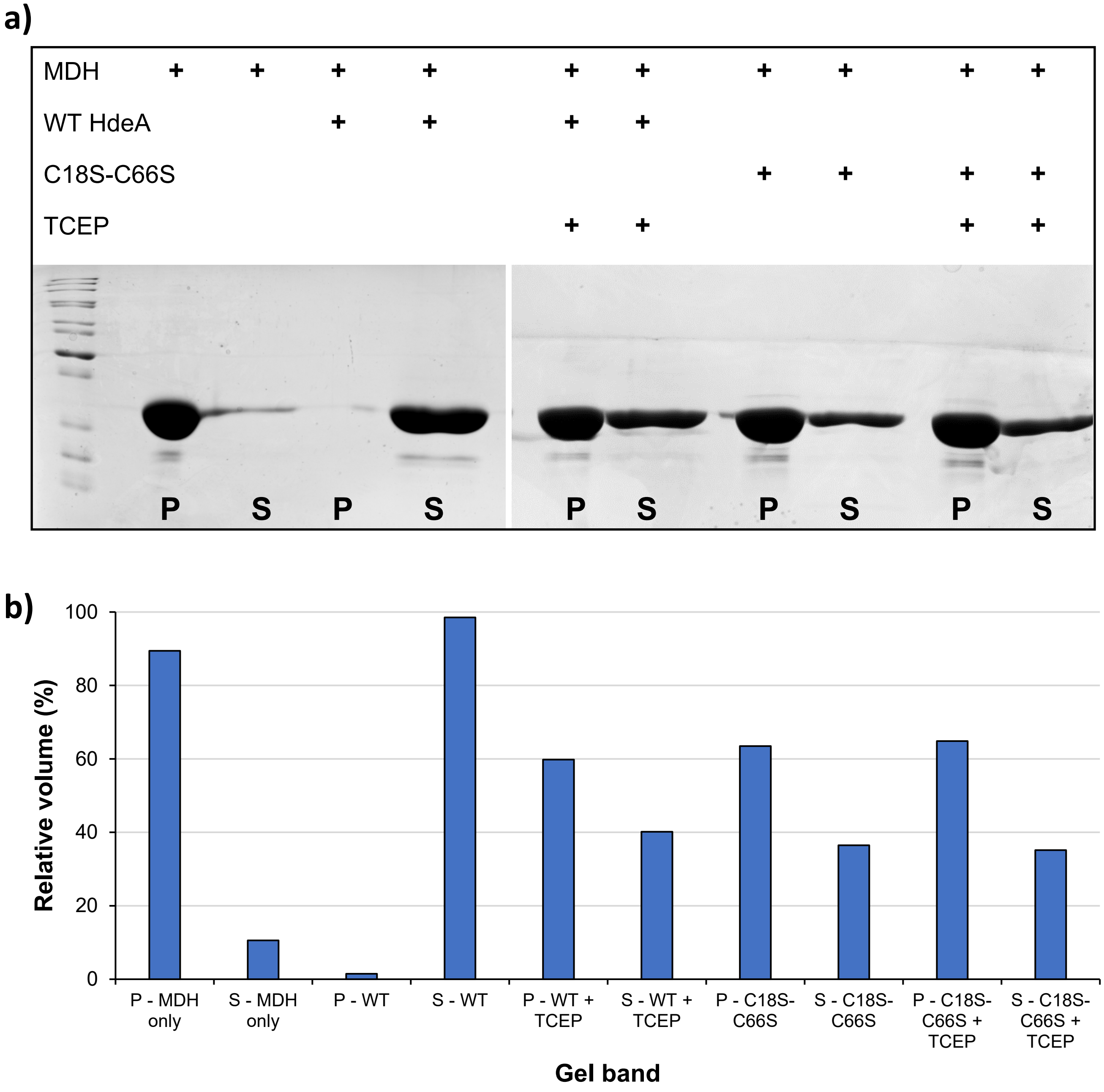


**Figure S6. a)** SDS-PAGE gels showing the HdeA chaperone activity assay at pH 2.0. “P” and “S” labels represent lanes containing pellet (aggregated MDH) or supernatant (soluble MDH), respectively. “+” signs in the table above the gels indicate the presence of the indicated component. Gel images have been adjusted for brightness and contrast, but the adjustment has not eliminated or obscured any bands. **b)** Plot of volume integration data from Bio-Rad Image Lab software. Values correspond to % relative volume (each pellet-supernatant pair adds up to 100%).
